# Retrieved Sequence Augmentation for Protein Representation Learning

**DOI:** 10.1101/2023.02.22.529597

**Authors:** Chang Ma, Haiteng Zhao, Lin Zheng, Jiayi Xin, Qintong Li, Lijun Wu, Zhihong Deng, Yang Lu, Qi Liu, Lingpeng Kong

## Abstract

The advancement of protein representation learning has been significantly influenced by the remarkable progress in language models. Accordingly, protein language models perform inference from individual sequences, thereby limiting their capacity to incorporate evolutionary knowledge present in sequence variations. Existing solutions, which rely on Multiple Sequence Alignments (MSA), suffer from substantial computational overhead and suboptimal generalization performance for de novo proteins. In light of these problems, we introduce a novel paradigm called Retrieved Sequence Augmentation (RSA) that enhances protein representation learning without necessitating additional alignment or preprocessing. RSA associates query protein sequences with a collection of structurally or functionally similar sequences in the database and integrates them for subsequent predictions. We demonstrate that protein language models benefit from retrieval enhancement in both structural and property prediction tasks, achieving a 5% improvement over MSA Transformer on average while being 373 times faster. Furthermore, our model exhibits superior transferability to new protein domains and outperforms MSA Transformer in de novo protein prediction. This study fills a much-encountered gap in protein prediction and brings us a step closer to demystifying the domain knowledge needed to understand protein sequences. Code is available at https://github.com/HKUNLP/RSA.

## 1 Introduction

Proteins, as fundamental yet complex components of life, exhibit a diverse range of functions within organisms [17]. The enigmatic nature of these macromolecules originates from the intricate interplay between their sequences, structures, and functions, which is influenced by both physics and evolution [47]. Commonly, language models are employed to model protein sequences by generating amino acid distributions that align with co-occurrence probabilities observed in nature, thus encapsulating structural and evolutionary information within representations. While this approach has proven effective [14, 30, 35, 38, 45], extracting adequate information from the sequence alone can be inadequate in addressing challenges in capturing fine-grained evolutionary knowledge [26, 35].

Retrieval-augmented models have demonstrated efficacy in incorporating domain knowledge and improving intricate reasoning abilities in natural language processing and machine learning [18, 20, 32, 52]. These models employ a large-scale memory of sequences as a knowledge base and utilize multiple related input sequences instead of a single input to establish connections with implicit knowledge. Such approach offers the potential for more interpretable and modular knowledge capture [19], enabling rapid generalization to novel domains [32, 8]. To this end, we investigate the enhancement of protein language model predictions through a straightforward retrieval-based augmentation that perform prediction based on related sequences. This process is particularly related with evolution, as sequentially similar proteins, or homologs, often result from evolutionary selection and are more likely to possess shared functional and structural characteristics with the target protein [4, 12, 56].

In this work, we explore **R**etrieved **S**equence **A**ugmentation (RSA) method as a general framework for augmenting protein representations. Specifically, RSA employs a pre-trained dense sequence retriever to retrieve protein sequences that are similar to the query sequence both in terms of homology as well as structure. By learning these sequences alongside the original input, the model incorporates external knowledge and transfers it to new domains. Our assessment of this method consists of comprehensive experiments conducted across seven distinct tasks, encompassing protein structure, function, evolution, and engineering, which require diverse knowledge. Employing a vast database of approximately 40 million protein sequences, we show that a retrieval-based approach leveraging this data consistently outperforms state-of-the-art methods.

In recent studies [30, 43, 55, 29], models such as MSA Transformer and AlphaFold2 have demonstrated remarkable success utilizing Multiple Sequence Alignment (MSA) of protein homologs as input features. However, these models heavily depend on the MSA alignment process, which is thought to underscore co-evolutionary features. This procedure is notoriously computationally intensive - for instance, it takes HHblits [44] 10 seconds for a single iteration search on Pfam using 64 CPUs. In contrast, RSA employs retrieved sequences from dense retrievers without requiring an alignment process, resulting in a 373-fold speed-up and on-the-fly processing, as shown in Figure 1. Additionally, RSA without additional pretraining outperforms a pre-trained MSA Transformer in downstream tasks, particularly for denovo proteins with few or no MSAs. Furthermore, we employ probabilistic analysis to integrate MSA-based approaches into the retrieval-augmentation paradigm and challenge conventional wisdom by demonstrating that, although residue alignment adheres to biological principles, modern language models do not solely depend on alignment to extract evolutionary information. Consequently, we conclude that retrieval augmentation for proteins as a general framework can be a sound replacement for MSA in terms of expressiveness, speed, and augmentation performance. Our major contributions can be summarized as:

**Figure 1:**
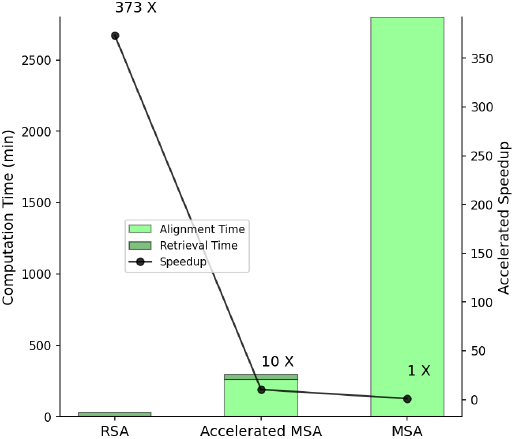
Illustration of speed up by RSA retrieval compared to MSA on secondary structure prediction dataset with 8678 sequences. Accelerated MSA refers to the MSA Transformer with MSA sequences retrieved by our RSA retriever.

**Figure 2:**
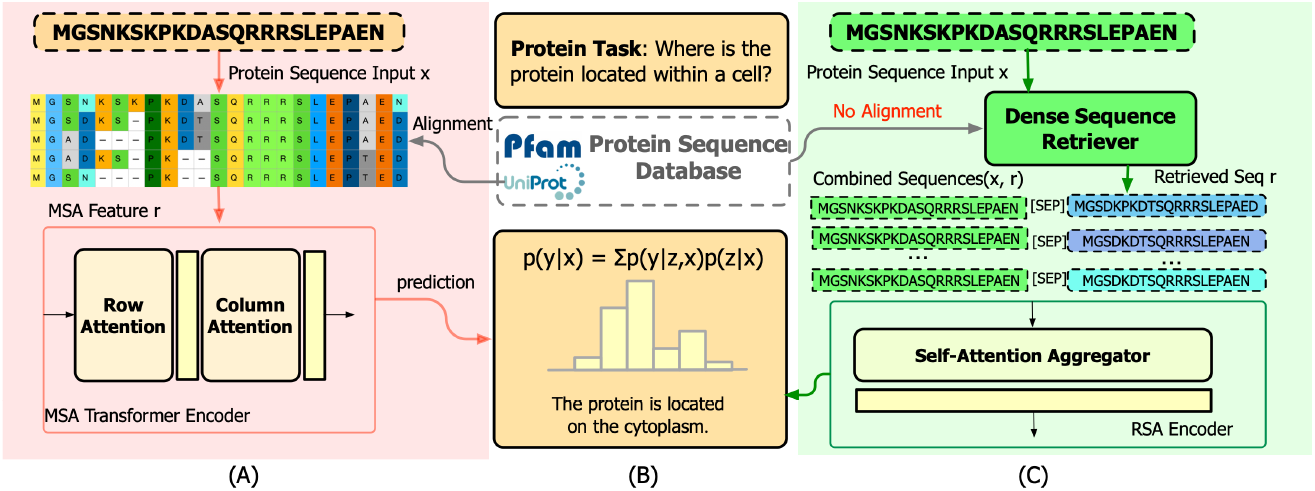
Comparison between MSA Transformer and RSA. (A) MSA Transformer aligns query to the protein database and use axial attention to encode MSA feature. (B) Overview of the probablistic framework. (C) RSA initially extracts related protein sequence from the database with a dense retriever. Subsequently, the query protein is augmented into pairs with each retrieved datum, which are input into the protein model for relevant tasks. Both MSA Transformer and RSA fall within the retrieval framework; however, RSA doesn’t require the alignment process.

- The novel investigation of retrieval-augmented protein language models and the proposal of the first alignment-free, efficient framework, RSA, for enhancing protein representations.
- Our theoretical establishment of a unified framework reveal two significant insights: (1) MSA-augmented methods are essentially retrieval-augmented language models. Their performance can be explained by the injection of evolutionary knowledge. (2) The *O*(*N* ^2^) complex alignment process is less necessary for deep protein language models.
- The demonstration that pre-trained dense retrievers offer greater efficiency and competitive efficacy in extracting homologous sequences and structurally similar sequences.

## 2 Related Work

### Retrieval-Augmented Language Models

The integration of non-parametric memory retrieval and parametric models has been an intriguing approach for many problems [31, 19, 21, 10, 59]. Retrieval-augmented language models explicitly introduce related knowledge from the memory and have shown improved performance in complex reasoning [50] and generalization [32, 8] to new domains. Our RSA method is motivated by retrieval-augmented language models [19, 21], though we specifically focus on injecting protein knowledge and adapt the model for token-level tasks and better efficiency.

### Protein Language Model

To model and further understand the protein sequence data, language models are introduced to train on mass data [23, 1]. Large scale pre-training enables language models to learn structural and evolutionary knowledge [14, 30, 35]. Despite these successes, many important applications still require MSAs and other external knowledge [43, 30, 22, 61, 29, 42]. MSAs have been shown effective in improving representation learning, despite being extremely slow and costly in computation. Hu et al. [26] and Hong et al. [24] use dense retrieval to accelerate multiple sequence augmentation, while still dependent on alignment procedures. Recent work [16, 35, 53, 11] explores MSA-free language models though additional pre-training is involved. We take this step further to investigate retrieval-augmented protein language models.

## 3 Augmenting Protein Representations with Retrieved Sequences

In this section, we introduce a unified probabilistic framework to connect the MSA-based models with retrieved augmentations. This framework offers a novel holistic view on understanding these models, that is the retrieved protein sequences enhance the performance of pre-trained protein models by providing evolutionary knowledge in a similar way as MSA sequences do. Furthermore, we emphasize design elements that inspire our methodology for achieving increased efficiency and adaptability.

### 3.1 Background and Problem Statement

Given a protein *x* = [*o*_1_, *o*_2_, …*o*_*L*_] comprising of *L* amino acids, the objective of a protein language model is to learn an embedding transferable to subsequent tasks. The embedding, represented as Embed(*x*) = [*h*_1_, *h*_2_, …*h*_*L*_], where *h*_*i*_ ∈ ℝ^*d*^ denotes a *d*-dimensional token representation for *o*_*i*_. The aim is to learn *p*(*y*|*x*) for predicting the properties of the sequence. For token property prediction tasks (e.g., secondary structure prediction) and pairwise prediction tasks (e.g., contact prediction), a prediction should be allocated to each token/pair, i.e., *p*(*y*_*i*_|*o*_*i*_) or *p*(*y*_*ij*_|*o*_*i*_, *o*_*j*_).

One way to construct an evolution-informed representation involves encoding MSA input into corresponding representations. We consider MSAs as *N* aligned protein homologs *r*_1_, …, *r*_*N*_. Prior studies [55, 29] encode MSA as co-evolution statistics features *R*_1…*N*_ and aggregate these features to derive the representation, while MSA Transformer [43, 30] perceives MSA as a matrix, employing axial attention to extract salient evolutionary traits. Here we also denote retrieved sequences as *r*_1_, …, *r*_*N*_ and their features as *R*_1…*N*_, though no alignment is performed to these sequences.

### 3.2 Protein Retrieval Augmentations: A Unified Framework

Inspired by Guu et al. [19], *protein retrieval augmentation*, that aims to unify several state-of-the-art evolution augmentation methods. Specifically, we consider these methods as learning a downstream predictor *p*(*y*|*x*) based on an aggregation of homologous protein representations *R*_1…*N*_. From the view of retrieval, *p*(*y*|*x*) is decomposed into two steps: *retrieve* and *predict*. For a given input *x*, the retrieve step first finds possibly helpful protein sequence *r* from a sequence corpus ℛ and then predict the output *y* conditioning on this retrieved sequence. We treat *r* as a latent variable and in practice, we approximately marginalized it out with top-*N* retrieved sequences:

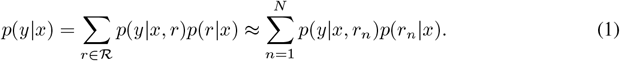

The probability *p*(*r*|*x*) denotes the possibility that *r* is sampled from the retriever given *x*. Intuitively it measures the similarity between the two sequences *r* and *x*. This framework also applies to the MSA-based augmentation methods. We explain in detail using a state-of-the-art MSA-augmentation model *MSA Transformer* [43] as an example. In MSA Transformer, the layers calculate self-attention both row-wise and column-wise. Column-wise attention is defined as follows, given *W*_*Q*_, *W*_*K*_, *W*_*V*_, *W*_*O*_ as the parameters in a typical attention function:

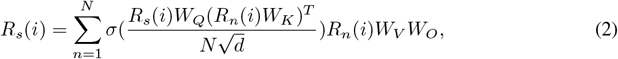

where *R*_*n*_(*i*) denotes the *i*-th token representation of the *n*-th MSA sequence after performing the row-wise attention. Note that in MSA input, the first sequence *r*_1_ is defined as the original sequence *x*. Then for a token prediction task, we define the *i*-th position output as *y* and the predicted distribution *p*(*y*|*x*) can be expressed as:

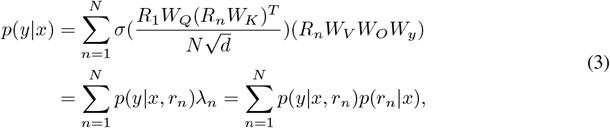

Where 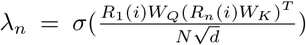 is the weighting norm that represents the similarity of retrieved sequence *r*_*n*_ and original sequence *x*; *p*(*y*|*x, r*_*n*_) is a predictor that maps the row-attention representation of *r*_*n*_ and *x* to label.

Eq.3 gives a retrieval-augmentation view of MSA Transformer that essentially retrieves homologous sequences with multiple sequence alignment and aggregates representations of homologous sequences with regard to their sequence similarity. Taking one step further, we define a set of design dimensions to characterize the retrieving and aggregation processes. We detail the design dimensions below and illustrate how popular models (§D) and our proposed methods (§4) fall along them in Table 1.

**Table 1:** Protein Retrieval Augmentation methods decomposed along a different axis. We formulate the aggregation function in the sequence classification setting and use a feed-forward neural network FFN(·) to map representations to logits. The proposed variants vary in design axis from the existing methods. ^†^Note that MSA Transformer performs the aggregation in each layer of axial attention, which differs from other variants.

| Method | Retriever Form | Alignment Form | Weight λ<sub>n</sub> | Aggregation Function |
|---|---|---|---|---|
| **Existing Methods** |  |  |  |  |
| Potts Model | MSA | Aligned | — | — |
| Co-evolution Aggregator | MSA | Aligned | $\frac{1}{N}$ | FFN(∑<sub>i=1</sub><sup>N</sup> R<sub>n</sub>(i)λ<sub>n</sub>) |
| MSA Transformer | MSA | Aligned | $\sigma(\frac{XW_Q(R_nW_K)^T}{N\sqrt{d}})$ | FFN(∑<sub>i=1</sub><sup>N</sup> R<sub>n</sub>(i)λ<sub>n</sub>)<sup>†</sup> |
| **Proposed Variants** |  |  |  |  |
| Unaligned MSA Augmentation | MSA | Not Aligned | σ(−\|\|X − R<sub>L</sub>\|\|<sub>2</sub>) | ∑<sub>i=1</sub><sup>N</sup> FNN(R<sub>n</sub>(i))λ<sub>n</sub> |
| Accelerated MSA Transformer | Dense Retrieval | Aligned | $\sigma(\frac{XW_Q(R_nW_K)^T}{\lambda(N,d)})$ | FFN(∑<sub>i=1</sub><sup>N</sup> R<sub>n</sub>(i)λ<sub>n</sub>) |
| Retrieval Sequence Augmentation | Dense Retrieval | Not Aligned | σ(−\|\|X − R<sub>n</sub>\|\|<sub>2</sub>) | ∑<sub>i=1</sub><sup>N</sup> FNN(Embed(x; r<sub>n</sub>))λ<sub>n</sub> |

- **Retriever Form** indicates the retriever type used. Multiple Sequence Alignment is a discrete retrieval method that uses alignment [58] to find homologous sequences. Dense retrieval [27] has been introduced to accelerate discrete sequence retrieval.
- **Alignment Form** indicates whether retrieved sequences are aligned.
- **Weight Form** is the aggregation weight of homologous sequences, as the *p*(*r*_*n*_|*x*) in Eq. 3. Here we denote this weight as *λ*_*n*_. Traditionally, aggregation methods consider different homologous sequences to be similarly important and use average weighting. MSA Transformer uses a weighted pooling method though the weights of *λ*_*n*_ use global attention and are dependent on all homologous sequences.
- **Aggregation Function** is how the representations of homologous sequences are aggregated to the original sequence to form downstream prediction, as in *p*(*y*|*x, r*). For example, considering the sequence classification problem, a fully connected layer maps representations to logits. The retrieval augmentation probabilistic form first maps each representation to logits *p*(*y*|*x, r*_*n*_) and then linearly weight the logits with *λ*_*n*_ in Eq. 3.

Our discussion and formulation so far reach the conclusion that retrieval augmentation serves as a comprehensive framework capable of extracting evolutionary knowledge, akin to multiple sequence alignment (MSA) augmentation methods. This underlines the prospects of retrieval sequence alignment (RSA) superseding MSA augmentations as an efficient and generalizable approach.

However, MSA-based methods claim a few advantages: the *alignment* process can help the model capture column-wise residue evolution; and the *MSA Retriever* uses a discrete, token-wise search criterion that ensures all retrieved sequences are homology. We propose two novel variants to help verify these claims: 1) **Unaligned MSA Augmentation** uses the homologous sequences from MSA to augment representations without alignment and 2) **Accelerated MSA Transformer** explores substituting the discrete retrieval process in MSA with a dense retriever. An empirical study of the performance of these models can be found in §5.6.

## 4 Our Approach

Existing knowledge augmentation methods for protein representation learning are either designed for a specific task or require cumbersome data preprocessing. Motivated by the potential of pre-trained retrievers to identify proteins that are homologous or geometric similar, we propose a pipeline, RSA (**R**etrieval **S**equence **A**ugmentation), to directly augment protein models on-the-fly. RSA follows the *retrieve-then-predict* framework in Eq. 1. It comprises of a neural sequence retriever *p*(*r*|*x*), and a protein model that combines both original input and retrieved sequence to obtain prediction *p*(*y*|*x, r*).

### RSA Retriever

The retriever is defined as finding the sequences that are semantically close to the query. Denote retriever model as *G* which encode protein sequence and output embeddings.

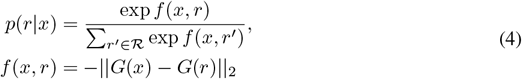

The similarity score *f* (*x, r*) is defined as the negative L2 distance between the embedding of the two sequences. The distribution is the softmax distribution over similarity scores.

For protein retrieval, we aim to retrieve protein sequences that have similar structures or are homologous to the query sequence. Motivated by the k-nearest neighbor retrieval experiment with ESM-1b

[45] pre-trained embeddings (as shown in Table 2), we implement the embedding functions using a 34-layer ESM-1b encoder. We obtain sequence embeddings by performing average pooling over token embeddings. Note that finding the most similar proteins from a large-scale sequence database is computationally heavy. To accelerate retrieval, we use Faiss indexing [28], which uses clustering of dense vectors and quantization to allow efficient similarity search at a massive scale.

**Table 2:** Recall and Precision for retrieving top 100 protein sequences with ESM1b embeddings. In dataset Pfam and SCOPe, we test whether retrieved proteins are of the same Family, Superfamily, or Fold as query protein.

| Retrieval Task | Type | Recall | Precision |
|---|---|---|---|
| Pfam - Family | Homology | 100 | 90.42 |
| SCOPe - Fold | Structural | 100 | 65.98 |
| SCOPe - Superfamily | Structural | 100 | 46.00 |
| SCOPe - Family | Structural | 100 | 24.71 |

### Retrieval Augmented Protein Encoder

Given a sequence *x* and a retrieved sequence *r* with length *L* and *M* respectively, the protein encoder combines *x* and *r* for prediction *p*(*y*|*x, r*). To make our model applicable to any protein learning task, we need to augment both sequence-level representation and token-level representation. To achieve this, we concatenate the two sequences before input into the transformer encoder, which uses self-attention to aggregate global information from the retrieved sequence *r* into each token representation.

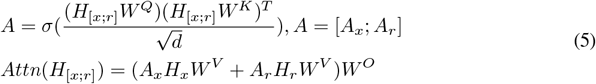

where 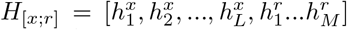 denotes the input embedding of original and retrieved sequences. The output token representation *h*_*i*_ automatically learns to select and combine the representation of retrieved tokens. This can also be considered a soft version of MSA alignment. After computing for each pair of (*x, r*), we aggregate them by weight *p*(*r*|*x*) defined in Eq. 4.

### Training

For downstream finetuning, we maximize *p*(*y*|*x*) by performing training on the retrieval augmented protein encoder. We freeze the retriever parameters during training. For a query sequence of length *L* with *N* retrieved proteins, suppose the length of retrieved proteins *L*^′^ ≤ *L* the computation cost is *N* times the original model, *O*(*NL*^2^) for a transformer encoder layer, which is as efficient as MSA Transformer with a *O*(*NL*^2^) + *O*(*N* ^2^*L*) computation cost.

## 5 Experiments

In this section, we conduct comprehensive experiments on 7 tasks to answer the following three questions: (1) Does RSA enhance protein representation learning concerning downstream performance, generalizability, and efficiency? (2) Is the alignment process in MSA dispensable? (3) What knowledge do the retrieved sequences provide to enhance the accuracy of downstream predictions? Experimental setup are briefly explained in §5.1 with more information in the appendix.

### 5.1 General Setup

#### Downstream Task

In order to evaluate the performance of our trained model, seven tasks are introduced, namely secondary structure prediction [33], contact prediction [3], remote homology prediction [25], subcellular localization prediction [2], stability prediction [46], protein-protein interaction [39] and structure prediction on CASP14 [34]. Please refer to Appendix Table 9 for more statistics of the datasets. The train-eval-test splits follow TAPE benchmark [41] for the first four tasks and PEER benchmark [54] for subcellular localization and protein-protein interaction.

#### Retriever and MSA Setup

Limited by available computation resources, we build a database on Pfam [13] sequences, which covers 77.2% of the UniProtKB [5] database and reaches the evolutionary scale. We generate ESM-1b pre-trained representations of 44 million sequences from Pfam-A and use Faiss [27] to build the retrieval index. For a fair comparison, the MSA datasets are also built on the Pfam database. We use HHblits [44] to extract MSA, searching for 3 rounds with threshold 1e-3.

#### Baselines

We apply our retrieval method to both pre-trained and randomly initialized language models. Following Rao et al. [41] and Rao et al. [43], we compare our model with vanilla protein representation models, including LSTM[36], Transformers[51] and pre-trained models ESM-1b[45], ProtBERT[15]. We also compare with state-of-the-art knowledge-augmentation models: Potts Model[6], MSA Transformer[43] that inject evolutionary knowledge through MSA, OntoProtein[62] that uses gene ontology knowledge graph to augment protein representations and PMLM[22] that uses pair-wise pretraining to improve co-evolution awareness.

#### Training and Evaluation

To demonstrate RSA as a general method, we perform experiments both with a shallow transformer encoder, and a large pre-trained ProtBERT encoder. The Transformer model has 512 dimensions and 6 layers. Also, we combined our method with popular pre-trained protein folding architectures ESMFold and AlphaFold2. All self-reported models use the same truncation strategy and perform parameter searches on the learning rate, warm-up rate, and batch size.

### 5.2 Main Results

We show the result for downstream tasks in Table 3, including models with/without pretraining, and with/without knowledge augmentations. We form the following conclusion: **Retrieval Sequence Augmentations perform on par with or even better than other knowledge-augmented methods without additional pre-training**. The last two blocks compare our method with previous augmentation methods. Our method outperforms MSA Transformer on average by 5% and performs on par with PMLM on structure and evolution prediction tasks. Notably, both MSA Transformer and PMLM perform additional pre-training with augmentations, while our method uses no additional pre-training. From the results, we can see that RSA combined transformer model also improves by 10% than other shallow models, demonstrating the effectiveness of our augmentation to both shallow models and pre-trained models. We also study retrieval sequence augmentations on pre-trained protein folding models in Table 4. Despite RSA was implemented without additional fine-tuning on folding models, we achieve a 2% improvement both on ESMFold and AlphaFold2.

**Table 3:** Main Results for vanilla protein representation learning methods, knowledge-augmented baselines and our proposed RSA method. Note that *italized* result is reported by corresponding related work. The last column reports average result on all six tasks. For MSA Transformer and RSA, we all use 16 sequences (N=16) for augmentation. For Gremlin Potts model, we use the full MSA.

| Method | Pretrain | Knowledge Pretrain | Knowledge Injection | SSP | Contact | Homology | Stability | Loc | PPI | Avg |
|---|---|---|---|---|---|---|---|---|---|---|
| Transformer | × | × | × | 0.384 | 0.274 | 0.101 | 0.422 | 0.541 | 0.616 | 0.345 |
| LSTM | × | × | × | *0.596* | *0.263* | *0.181* | *0.591* | *0.629* | *0.638* | 0.404 |
| RSA (Transformer backbone) | × | × | ✓ | 0.541 | 0.332 | 0.346 | 0.602 | 0.591 | 0.700 | 0.518 |
| ESM-1b | ✓ | × | × | *0.716* | *0.458* | *0.978* | *0.695* | *0.781* | *0.782* | 0.668 |
| ProtBERT | ✓ | × | × | 0.691 | 0.556 | 0.528 | 0.651 | 0.771 | 0.688 | 0.579 |
| MSA Transformer (MSA N=1) | ✓ | ✓ | × | 0.594 | 0.397 | 0.880 | 0.767 | 0.668 | 0.633 | 0.592 |
| Gremlin [6] | × | × | ✓ | — | 0.507 | — | — | — | — | — |
| MSA Transformer | ✓ | ✓ | ✓ | 0.654 | 0.618 | 0.958 | **0.796** | 0.694 | 0.751 | 0.672 |
| OntoProtein [62] | ✓ | × | ✓ | *0.68* | *0.40* | 0.96 | *0.75* | — | — | — |
| PMLM [22] | ✓ | ✓ | × | ***0.728*** | ***0.717*** | *0.946* | — | — | — | — |
| RSA (ProtBERT backbone) | ✓ | × | ✓ | 0.691 | **0.717** | **0.987** | 0.778 | **0.795** | **0.827** | **0.723** |

**Table 4:** Results for Structure Prediction on CASP14. AlphaFold-Acc uses accelerated-MSA and Percentage stands for the percentage of samples more precise than baseline prediction.

| Methods | TM-Score | Percentage |
|---|---|---|
| ESMFold | 0.678 |  |
| AlphaFold-single | 0.335 |  |
| ESMFold-RSA | 0.693 | 27.7% |
| AlphaFold-RSA | 0.359 | 45.5% |
| AlphaFold-Full | **0.747** |  |
| AlphaFold-Acc | 0.551 | 19.7% |

### 5.3 Retrieval Augmentation for Domain Adaptation

We investigate the model’s transfer performance in domains with distribution shifts. We train our model on the Remote Homology dataset, and test it on three testsets with increasing domain gaps: proteins within the same Family, Superfam, and Fold as the training set respectively. The results are in Table 5. Our model surpasses MSA Transformer by a large margin on shifted domains, especially from 0.5032 to 0.6770 on Superfamily. This proves our models to be more reliable for domain shifts, illustrating that retrieval facilitates the transfer across domains.

**Table 5:**
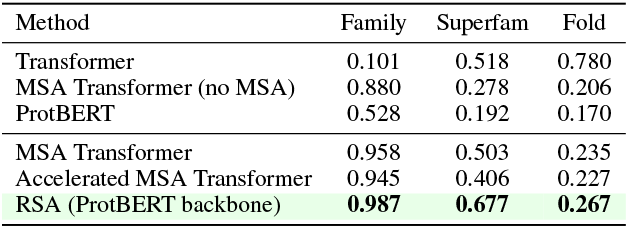
The table shows remote homology prediction performance with increasing domain gaps: Family, Superfamily and Fold.

| Method | Family | Superfam | Fold |
|---|---|---|---|
| Transformer | 0.101 | 0.518 | 0.780 |
| MSA Transformer (no MSA) | 0.880 | 0.278 | 0.206 |
| ProtBERT | 0.528 | 0.192 | 0.170 |
| MSA Transformer | 0.958 | 0.503 | 0.235 |
| Accelerated MSA Transformer | 0.945 | 0.406 | 0.227 |
| RSA (ProtBERT backbone) | **0.987** | **0.677** | **0.267** |

Furthermore, we test our model on a challenging problem in protein prediction, the prediction for proteins with few homologs, i.e. de novo (synthesized) proteins and orphan proteins [16, 53]. This task is especially difficult for MSA-based methods as alignment-based method often fails to generate MSA for these proteins, resulting in degraded performance. We test our model on 108 De Novo proteins from PDB [9] for the contact prediction task. It can be seen in Figure 3 that, RSA exceeds MSA transformer on 63.8% of data, demonstrating that RSA is more capable of locating augmentations for out-of-distribution proteins. We also test our model on the structure prediction task with 16 targets from CASP14-FM. CASP14-FM are considered more difficult because the absence of related templates requires the prediction methods to rely on de novo modeling techniques. We compare RSA augmented ESMFold and AlphaFold2 model with baselines in Figure 3, showing improved or competitive prediction on the majority of the targets. The results also show that our model surpasses MSA-based methods in transferring to unseen domains.

**Figure 3:**
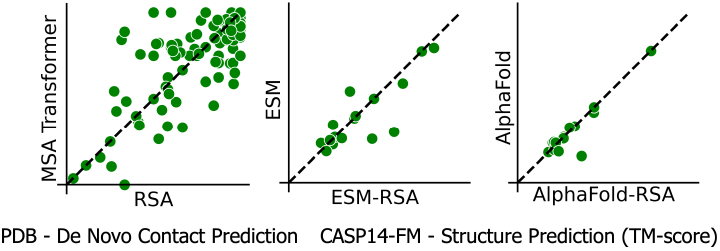
Prediction on proteins with few homologs, including contact prediction result on PDB de novo proteins and structure prediction result on CASP14-FM.

**Figure 4:**
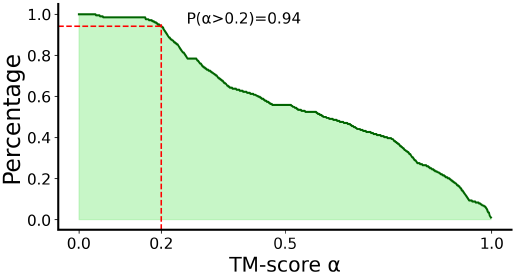
The cumulative distribution of TM-scores for proteins from dense retrieval. The value at *α* shows the probability that TM-score is larger than *α*.

### 5.4 Retrieval Speed

A severe speed bottleneck limits the use of previous MSA-based methods. In this part, we compare the computation time of RSA with MSA and an accelerated version of MSA as introduced in § 3.2. As shown in Figure 1, alignment time cost is much more intense than retrieval time. Even after reducing the alignment database size to 500, accelerated MSA still need 270 min to build MSA. At the same time RSA only uses dense retrieval, and is accelerated 373 times. Note that with extensive search, MSA can find *all* available alignments in a database. However, this would be less beneficial to deep protein language models as the memory limit only suffices a few dozens of retrieved sequences.

Also, MSA is limited by its cubersome construction of retrieval HHM profile to perform HHM-HHM search. Previous work mentioned that it may take many days to construct a custom HHblits database for a large database, though no precise time is given [60].By contrast, RSA only needs to build the pre-trained features for the database, which can be accelerated with GPUs and batch forwarding. We build the retrieval index for Pfam with 16 Tesla V100 GPUs in 20 hours.

### 5.5 Ablation Study

#### Ablation on Retriever: Unaligned MSA Augmentation

We ablate RSA retriever by using MSA retrieved proteins as augmentations to our model, denoted as Unaligned MSA Augmentation. As shown in Table 6, Unaligned MSA Augmentation performs worse than our RSA model, especially on the Stability dataset, where the performance drops from 0.778 to 0.7443. It thus confirms the ability of our dense retriever to provide more abundant knowledge for protein models.

**Table 6:**
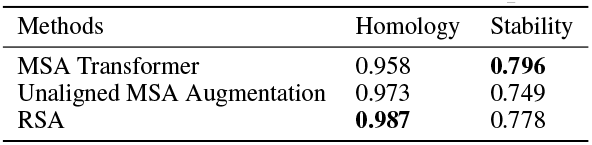
Results for MSA Transformer and Unaligned MSA Augmentation on Homology and Stability task. Both models use MSA as inputs, but Unaligned MSA Augmentation unaligns MSA and augments the model by concatenating MSA sequence to the input.

| Methods | Homology | Stability |
|---|---|---|
| MSA Transformer | 0.958 | **0.796** |
| Unaligned MSA Augmentation | 0.973 | 0.749 |
| RSA | **0.987** | 0.778 |

#### Ablation on Retriever: Ablation on Retrieval Number

Our study examines the effect of injected knowledge quantity for RSA and all retrieval baselines. The results are listed in Table 8. We select the Contact dataset because all baseline models are implemented on this dataset. RSA and all baselines perform consistently better as the retrieval number increases. Also, our model outperforms all baseline models for all augmentation numbers.

#### Ablation on Aggregation

We compare RSA with Accelerated MSA Transformer to evaluate whether our aggregation method is beneficial for learning protein representations. Note that only part of the retrieved sequences that satisfy homologous sequence criteria are selected and utilized during alignment. As shown in Table 7, the performance of the Accelerated MSA Transformer drops a lot compared to RSA. In contrast to MSA type aggregation, which is restricted by token alignment, our aggregation is more flexible and can accommodate proteins with variant knowledge.

**Table 7:** Results for MSA Transformer and Accelerated MSA Transformer on downstream tasks. Accelerated MSA Transformer uses MSA built from dense retrieval sequences.

| Tasks | MSA<br>Transformer | Accelerated MSA<br>Transformer | RSA |
|---|---|---|---|
| SSP | 0.654 | 0.634 | **0.691** |
| Contact | 0.618 | 0.608 | **0.717** |
| Homology | 0.958 | 0.945 | **0.987** |
| Stability | **0.796** | 0.767 | 0.778 |
| Loc | 0.694 | 0.682 | **0.795** |
| PPI | 0.751 | 0.679 | **0.827** |

### 5.6 Discussion on Alignment: Is It Necessary?

#### Does alignment makes better retriever?

Table 7 illustrates that Accelerated MSA Transformer performs near to MSA Transformer (MSA N=16) for most datasets, except for Stability and PPI on which our retriever failed to find enough homologous sequences, as Figure 5 demonstrates. Also, as shown in the figure, our dense retriever is capable of finding homologous sequences for most tasks and surpasses alignment in E-value.

**Figure 5.**
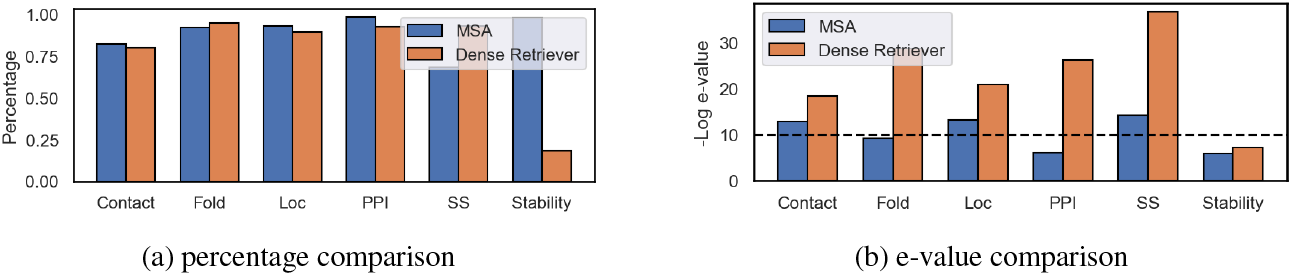
(a) Plot of the percentage of sequences that have found homologs on datasets for six tasks.(b) Plot of the -log(E-values) of MSA and Dense Retriever obtained sequences. E-values of both methods are obtained with HHblits[44]. Sequences with -log E-value >10 are high-quality homologs.

#### Is MSA alignment necessary for expressing knowledge?

To support that MSA alignment is not necessary, we compare Unaligned MSA Augmentation to the original MSA transformer. As revealed by the results in Table 6. Unaligned MSA Augmentation performs close to the MSA transformer. This confirms our declaration that self-attention is capable of extracting knowledge without alignment.

### 5.7 Retrieved Protein Interpretability

In this section, we give an intuitive analysis of what constitutes knowledge for protein understanding and why retrieved sequences can be used for improving protein reprensetations. We cover two major aspects of biology sequences, homology and geometry.

#### Dense Retrievers Find Homologous Sequences

In this part, we analyze whether retrieved sequences are homologous. As illustrated in Figure 5(a), across all six datasets, our dense retriever retrieved a high percentage of homologous proteins that can be aligned to the original protein sequence, comparable to traditional MSA retrievers. We additionally plot each dataset’s negative log E-values distribution in Figure 5(b). Accordingly, dense retrieval show high potential for finding homologous sequences, which explains the ability of RSA to capture evolutionary knowledge.

#### RSA Retriever Find Structurally Similar Protein

In this section, we analyze whether retrieved sequences are structurally similar. In Figure 4, we plot the TM scores between the RSA retrieved protein and the origin protein on ProteinNet [3] test set. Most of the retrieved proteins exceed the 0.2 criteria, which indicates structural similarity, and about half are above the 0.5 criteria, which indicates high quality. Accordingly, this indicates that the dense retrieval algorithm is capable of finding proteins with structural knowledge.

## 6 Discussions

In this paper, we introduce a simple yet effective method to enhance protein representation learning. We demonstrate RSA as a fast yet high-performing method that has the potential to replace MSA-based. The most notable limitation is that our method is dependent on high quality pre-trained embeddings and the abundance of protein sequences. We intend to further scale up our RSA method to larger protein databases and pre-train a retriever on abundant data in future work.

## Appendix

### A Limitations and Failed Case Analysis

One notable limitation of our method RSA is that it is highly dependent on high-quality pre-trained embeddings and the abundance of protein sequences. We found that our retriever tends to perform better in a database that has more protein sequences – that have not been screened by a clustering algorithm, like Uniclust30. This could be explained by our nearest neighbor retrieval technique which often requires more similar sequences for augmentation. We also found different patterns in retrieval sequences from MSAs. Our retriever tends to show polarized retrieval quality, either finding many evolutionary close sequences or failing to find any homologous sequences. We believe this is due to the imbalanced training of pre-trained embeddings on different protein families and hope to mitigate this issue with further training on retrieval datasets.

We report other failed cases here for a more thorough view of our proposed method:

- Directly applying Accelerated MSAs to MSA-based pre-trained models often shows about 2-3% decrease on downstream performance than using original MSAs. However, Accelerated MSAs are 10 times faster.
- The performance of RSA improves marginally with more sequences when *N >* 16. This is because we use the softmax distribution over L2 metrics to perform weighting, thereby assigning low weights to sequences further from the query.
- We found that in protein folding tasks, performing Average Pooling on ESMFold/AlphaFold shows worse zero-shot performance than Max Pooling with a scoring model. This is due to the misalignment of protein structures and simple weighting could result in averaging the structures of proteins with different angles of view.

### B Broader Impact

In this section, we discuss the broader impact of RSA in terms of protein representation learning, de novo protein understanding, as well as the potential application to large language models.

#### RSA for Protein Representation Learning

Developing efficient protein representation learning methods will significantly improve the ability to analyze complex protein structures, functions, and interactions. This would lead to a more comprehensive understanding of biological processes at the molecular level, consequently boosting advancements in the fields of bioinformatics and computational biology. In this paper, we propose RSA as an efficient and effective protein representation learning methods, which will spur the development of protein representation learning methods. Notably, our method requires no alignment methods. The traditional alignment process in MSA often requires mass CPU engines mostly available to academics. Our method on the other hand only requires a small memory GPU like 3090Ti and we will publicize our retrieval index, promoting democratic research in this field.

#### RSA for De Novo Protein Understanding

We have shown in our work that RSA could perform De Novo Protein Understanding. This is particularly important for drug repurposing and virtual screening tasks [40] for drug discovery. This can contribute to the development of personalized medicine by facilitating the identification of disease-specific protein biomarkers and selecting molecular cures for various diseases. However, de novo protein understanding often relies on newly-designed protein databases, which may include sensitive information about individuals, such as their genetic makeup, or violates intellectual property rights. Ensuring the privacy and security of this data is critical to prevent misuse and protect individual rights

#### RSA as Tool for Large Language Models

In addition to the potential impacts in the field of biology, our method could also improve the ability of Large Language Models in biological sequence understanding. Currently, large language models like ChatGPT show difficulty in understanding protein sequences. We showcase how RSA could improve this ability with the combination of retrieval and chain of thought. This application is valuable in education and training, as uses could rapidly learn about proteins through chat models, which help educate the next generation of researchers in bioinformatics, computational biology, and related fields. This will lead to a more skilled workforce in the life sciences.

### C A Brief Recap on Proteins

Proteins are the end products of the decoding process that starts with the information in cellular DNA. As workhorses of the cell, proteins compose structural and motor elements in the cell, and they serve as the catalysts for virtually every biochemical reaction that occurs in living things. This incredible array of functions derives from a startlingly simple code that specifies a hugely diverse set of structures. In fact, each gene in cellular DNA contains the code for a unique protein structure. Not only are these proteins assembled with different amino acid sequences, but they also are held together by different bonds and folded into a variety of three-dimensional structures. The folded shape, or conformation, depends directly on the linear amino acid sequence of the protein. In fact, this phenomenon is denoted as the *sequence-structure-function paradigm*. Here we will emphasize four key concepts in protein understanding.

#### 1. What are proteins made of ?

Amino acids. Within a protein, multiple amino acids are linked together by peptide bonds, thereby forming a long chain. There are 22 alpha-amino acids, from which proteins are composed. We model these amino acids in a similar way in NLP, as tokens. A tokenizer breaks the protein sequences into amino acid tokens that could be modeled by protein language models.

#### 2. Protein structures

There are four levels of structures in protein, as illustrated in Figure 6:

**Figure 6:**
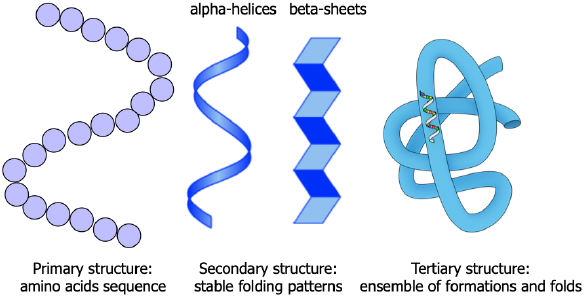
Illustrated explanation of protein levels of structures, primary structure, secondary structure and tertiary structure.

- Primary structure: amino acids sequence
- Secondary structure: stable folding patterns, including Alpha Helix, Beta Sheet.
- Tertiary structure: ensemble of formations and folds in a single linear chain of amino acids
- macromolecules with multiple polypeptide chains or subunits

Predicting protein structure is an important and difficult task. In this work, we also perform experiments on three tasks – secondary structure prediction, protein contact prediction (tertiary structure), and protein folding (tertiary structure), with increasing task difficulty.

#### 3. Protein Homology

Protein homology is defined as shared ancestry in the evolutionary history of life. There exists different kinds of homology, including orthologous homology that may be similar function proteins across species (human and mice *α*-goblin), and paralogous homology that is the result of mutations (human *α*-goblin and *β*-goblin). Homologies result in conservative parts in protein sequences, or leads to similar structures and functions.

#### 4. Multiple Sequence Alignments

A method used to determine conservative regions and find homologous sequences. An illustration (Figure 7) is given here to show how sequences are aligned. Aligned tokens may include the original amino acid, substitution, and deletions. The traditional way to generate MSA is using dynamic programming, with *O*(*L*^*N*^) complexity. Temporary methods use HMM-HMM alignment, as well as other acceleration methods. HH-Suite3 [48] reports a time complexity of *O*(*NL*^2^), which is still costly when performing alignment on a large database.

**Figure 7:**
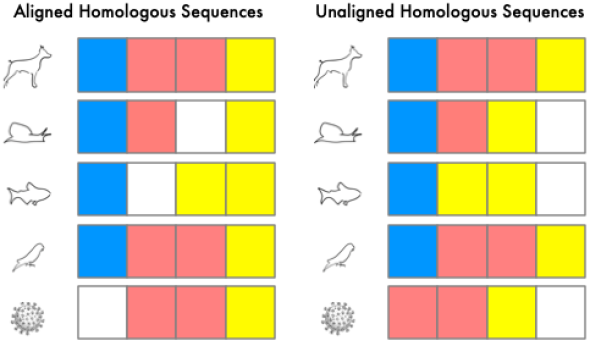
Illustrated difference of aligned and unaligned protein sequences. The white colour stands for the empty space in alignment “-”.

### D Overview of Previous Protein Representation Augmentation Methods

Below we introduce several state-of-the-art evolution augmentation methods for protein representation learning. These methods rely on MSA as input to extract representations. We use *x* to denote a target protein and its MSA containing *N* homologous proteins. We consider MSAs as *N* aligned protein homologs *r*_1_, …, *r*_*N*_. These studies [55, 29] encode MSA as co-evolution statistics features *R*_1…*N*_ and aggregate these features to derive the representation, while MSA Transformer [43, 30] perceives MSA as a matrix, employing axial attention to extract salient evolutionary traits. A unified view of these variants is available in Table 1 and §3.2 in the main paper.

#### Potts Model [6]

This line of research fits a Markov Random Field to the underlying MSA with likelihood maximization. This approach is different from other protein representation learning methods as it only learns a pairwise score for residues contact prediction. We will focus on other methods that augment protein representations that can be used for diverse downstream predictions.

#### Co-evolution Aggregator [55, 29]

One way to build an evolution informed representation is to use a MSA encoder to obtain the co-evolution related statistics. By applying MSA encoder on the *n*-th homologous protein in the MSA, we can get a total of *L* × *d* embeddings *R*_*n*_, each position is a *d* channel one-hot embedding indicating the amino acid type. We use *w*_*n*_ to denote the weight from *R*_*n*_ when computing the token representation *h*_*i*_:

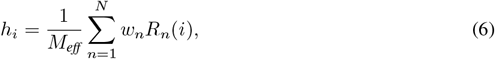

where 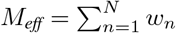 and 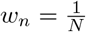. For contact prediction, pair co-evolution representation are computed in a similar way from the hadamard product:

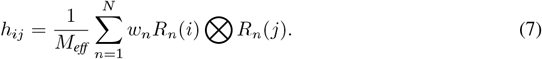

#### Ensembling Over MSA [42]

This approach aligns and ensembles representations of homologous sequences. Consider the encoder extract the same token representations for unaligned and aligned sequences. The ensembled token representation is:

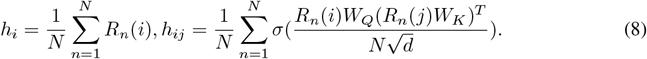

#### MSA Transformer [43]

In each transformer layer, a tied row attention encoder extracts the dense representation *R*_*n*_, then a column attention encoder

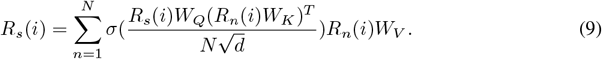

#### Knowledge Graph Augmentation [62, 63]

This line of research aims at incorporating factual knowledge in protein representations. Different from MSA-based methods that draw evolution knowledge from raw protein sequences, these methods are dependent on protein knowledge graphs that have been annotated by experts, therefore we only provide comparisons with these models in experimental studies and don’t incorporate them into our unified framework.

### E Experiment Setups

#### E.1 In-depth Introduction to Protein Tasks

##### Secondary structure prediction (SSP)

*Task Formulation*: 8-class classification *o*_*i*_ ↦ {0, 1, …, 7}

*Task Description*: Secondary structure prediction aims to predict the secondary structure of proteins, which indicates the local structures. This task predicts an 8-class label for each token, indicating which local structure this amino acid belongs to.

*Task Impact*: This task helps to determine whether a model captures protein local structure.

##### Contact prediction (Contact)

*Task Formulation*: 2-class classification (*o*_*i*_, *o*_*j*_) ↦ {0, 1}

*Task Description*: Contact prediction predicts the medium-range and long-range (distance >6) residue-residue contact, which measures the ability of models to capture global tertiary structures.

*Task Impact*: This task helps to determine whether a model captures protein tertiary structure. The assessment of this task focuses specifically on medium- and long-range interactions due to their crucial importance in the protein folding process.

##### Homology prediction (Homology)

*Task Formulation*: 1195-class classification *x* ↦ {0, 1 … 1194}

*Task Description*: Homology prediction aims to predict the fold label of any given protein, which indicates the evolutionary relationship of proteins.

*Task Impact*: Protein fold classification is important for both functional analysis and evaluating evolutionary knowledge.

##### Stability prediction (Stability)

*Task Formulation*: regression *x* ↦ ℝ

*Task Description*: Stability prediction is a protein engineering task, which measures the change in stability w.r.t. residue mutations.

*Task Impact*: Evaluate the ability of models to predict protein function as well as evaluate the ability of models to understand mutations, which is crucial for drug discovery and protein engineering.

##### Subcellular Localization (Loc)

*Task Formulation*: regression *x* ↦ {0, 1, …, 7}

*Task Description*: Subcellular localization refers to the process of determining the specific location or compartment within a cell where a particular molecule or protein resides. This information is essential for understanding the function and behavior of molecules or proteins, as their subcellular locations often dictate their roles in cellular processes, interactions with other molecules, and influence on cellular functions. For example, proteins on the cell membrane generally have signaling and regulatory functions.

*Task Impact*: This task is closely related to protein functions and roles in biological processes.

##### Protein-Protein Interaction (PPI)

*Task Formulation*: two-class classification (*x*_1_, *x*_2_) ↦ {0, 1}

*Task Description*: Protein-protein interaction predicts whether two proteins interact with each other.

*Task Impact*: This task is crucial for protein function understanding and drug discovery.

##### Protein Folding (Fold)

*Task Formulation*: *x* ↦ *S*, where *S* is the 3d-structure of protein, including all coordinates of atoms.

*Task Description*: Protein Folding predicts the structure of protein sequences.

*Task Impact*: This task is known to be challenging, and requires elaborated knowledge of protein local and global structure to make atomic predictions.

*Dataset Details*: We report test results on CASP14 public available targets. We also remove all sequences over 800 tokens due to the computation memory limit. The reported targets are: T1024, T1025, T1026, T1027, T1028, T1029, T1030, T1031, T1032, T1033, T1034, T1035, T1036s1, T1037, T1038, T1039, T1040, T1041, T1042, T1043, T1045s1, T1045s2, T1046s1, T1046s2,T1047s1, T1047s2, T1048, T1049, T1050, T1051, T1053, T1054, T1055, T1056, T1057, T1058, T1059, T1060s2, T1060s3, T1062, T1063, T1064, T1065s1, T1065s2, T1066s1, T1066s2, T1067, T1068, T1069s1, T1069s2, T1070, T1071, T1072s1, T1072s2, T1073, T1074, T1075, T1076, T1077, T1078, T1079, T1082, T1083, T1084, T1085, T1086, T1087, T1088, T1089, T1090, T1092, T1093, T1094, T1095, T1096, T1098, T1099, T1100, T1101. The blue targets are from CASP14-FM set. Table 9 gives the details of the datasets for these tasks.

##### De Novo Contact Prediction

We follow Chowdhury et al. [11] to curate a de novo dataset of 108 proteins from Protein Data Bank [Bank]. These proteins are originally designed de novo using computationally parametrized energy functions and are well-suited for out-of-domain tests. Note that different from orphan dataset, MSA can be built for this dataset, though showing a decline in quality.

#### E.2 HHblits Settings

For MSA datasets, We use HHblits [44] to perform alignment. The commands for MSA dataset construction is:

~~~
hhblits -cpu $CPU_NUM -i $INPUT_FILE -d $DATABASE_DIR -oa3m $OUTPUT_FILE -n
 1 -e 0.001
~~~

We also use HHblits to calculate E-value and determine whether we found homologous sequences in Figure 5 and §5.7 in the main paper. The commands for protein E-value calculation is:

~~~
hhalign -i query.fasta -d retrieved.fasta -o output.aln -e 0.001
~~~

#### E.3 Model Hyperparameters

All self-reported models use the same truncation strategy and perform parameter searches on the learning rate among [3*e* − 8, 3*e* − 6, 3*e* − 5, 3*e* − 4, 1*e* − 3], warm-up rate among [0, 0.08], seed among [111, 222, 333, 444, 555, 666], and batch size among [1, 2, 4, 8, 16]. For evaluation, we choose the best-performing model on the validation set and perform prediction on the test set. The best performing hyperparameters could be found in the file:

~~~
./RSA-code\scripts\$MODEL_NAME\run_$TASK_NAME.sh
~~~

Also, code with download instructions for dataset and retrieval index is available in the supplementary.

#### E.4 RSA and Variants Implementation Details

##### E.4.1 Retriever Implementation Details

First, we calculate the ESM-1b embeddings of the 44 million sequences in Pfam-A 32.0. We use 16 V100 GPUs to calculate the embeddings in a day. A GPU as small as 3090 Ti would be enough, though it would take longer. Then, we adopt Faiss [27] indexing to accelerate the retrieval process by clustering the pre-trained dense vectors. In our implementation, we use the Inverted file with Product Quantizer encoding Indexing and set the size of quantized vectors to 64, the number of centroids to 4096, and the number of probes to 8. The construction of the Faiss index takes roughly 30 minutes using 0.5% randomly selected protein embeddings for index training. All embeddings as well as their id are subsequently added to the index.

During retrieval, for each query sequence, we first use ESM-1b to calculate its embedding, and then using this embedding, we query faiss to find the top *N* nearest neighbor of this embedding, getting the distance and sequence id of retrieved sequences. L2 distances are used to measure sequence similarity.

##### E.4.2 ProtBERT-RSA Architecture and Implementation

Here we provide the details for ProtBERT-RSA Architecture. An illustration of this process is also available in Figure 8. Note that in Step 2 retrieval of Faiss index could be further accelerated with GPU. In Step 4, the predictions of pairwise augmentation could be accelerated with batching on GPU, concurrently predicting *k* augmented sequences at the same time.

**Figure 8:**
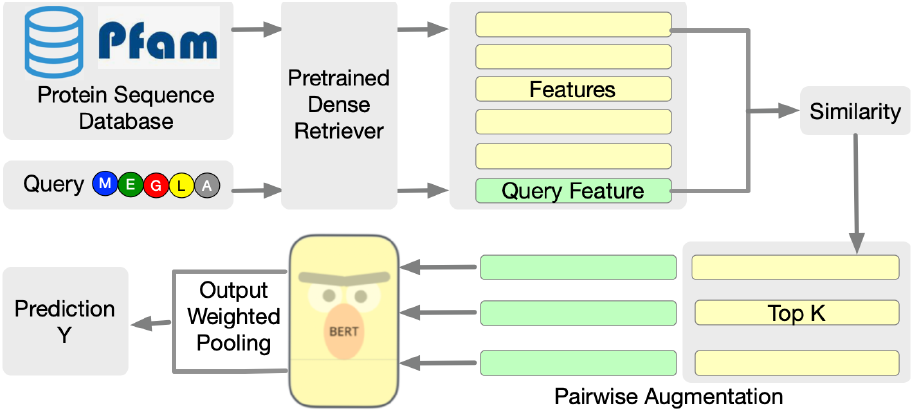
Detailed illustration of ProtBERT-RSA architecture.

However, for large pre-trained models and when *k* is very large, the batch computation may exceed memory limit. In this case, we provide implementation for gradient accumulation, which calculates loss and gradients for individual prediction (predictions_*i*_) and sum up the gradients with gradient accumulation. This implementation is a convex upperbound for the original loss function and we have validated its stability. This could also be implemented in batch size *n*, where each backward iteration calculates *k/n* retrieval augmentations, achieving trade-off between inference speed and memory limit.

~~~
Given query sequence $query, retrieval database $Faiss_Index, sequence
 database $Pfam, the number of retrieval $k, ProtBERT model $Model, and
 label $y.
Step 1. embedding = ESM_1b(query)
Step 2. distances, ids = Faiss_Index.retrieve(embedding, k)
retrieved_seqs = Pfam[ids]
Step 3. predictions_i = Model([query, retrieved_seq]), i=1,2,..k
Step 4. prediction = sum(predictions_i * softmax(distance_i))
Step 5. loss = loss_function(prediction, y), perform training
~~~

##### E.4.3 RSA for Protein Folding

The major difference of RSA prediction for protein folding from other tasks is that we use a ranker to choose the final prediction rather than using weighted pooling. This is due to the misalignment of protein structures and simple weighting could result in averaging the structures of proteins with different angles of view. We train the ranker together with pTM-score loss [35] and contrastive loss on a subset of 1000 randomly chosen proteins from Protein Data Bank. These proteins are distinct from CASP14 test set. The ranker takes in the original structure prediction of the protein sequence and the *k* augmented predictions, and generate the highest ranking prediction as the final result. As current protein folding models are very large, we only provide zero-shot testing results on these pre-trained models, without further finetuning on our pipeline.

~~~
Given query sequence $query, retrieval database $Faiss_Index, sequence
 database $Pfam, the number of retrieval $k, Folding model $Model,
 Ranking model $Ranker and label $y.
Step 1. embedding = ESM_1b(query)
Step 2. distances, ids = Faiss_Index.retrieve(embedding, k) retrieved_seqs = Pfam[ids]
Step 3. predictions_i = Model([query, retrieved_seq]), i=1,2,..k
Step 4. prediction = Ranker(predictions_i), i=1,2,..k
~~~

Due to the different model architectures of ESMFold and AlphaFold, we explain in details the inference pipeline of Model([query, retrieved]).

###### ESMFold-RSA

ESMFold is a **single** sequence protein folding model that consists of a protein representation model and a folding trunk based on the extracted representation. As illustrated in Figure 9(a), we concatenate query sequence with retrieved sequence and input them into the representation encoder. The encoder combines information from both query and retrieved sequence into query embedding via self-attention. Then we could use the pre-trained folding trunk to predict the structure of the query sequence. This pipeline could also be accelerated with batch prediction.

**Figure 9:**
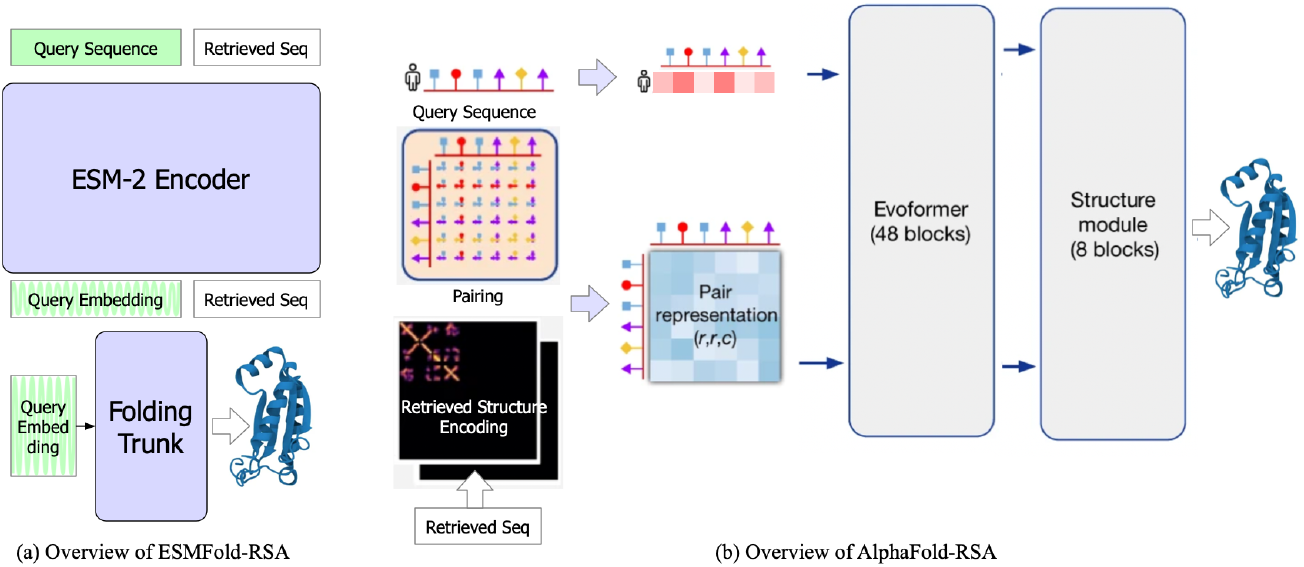
Illustration of the inference pipeline of RSA for Protein Folding

###### AlphaFold-RSA

Different from ESMFold, AlphaFold encoder takes both single sequence representation and pairwise representation as input. Therefore, as shown in FIgure 9(b), we generate the retrieved structure encoding with AlphaFold based on retrieved sequences, then we generate the structure of the query sequence based on the combination of single and pair representation. Note that we removed the template and MSA input in AlphaFold to ablation the effect of RSA.

##### E.4.4 Accelerated MSA

Accelerated MSA variant explores 165 substituting the discrete retrieval process in MSA with a dense retriever. We implement this method by first retrieving 500 sequences and then aligning these sequences with JackHMMer tool. Note that for most tasks we retrieve 500 sequences before alignment, as MSA Transformer can’t take in many sequences. The command for aligning is:

~~~
./jackhmmer -E 10.0 -A $aligned_file query.fasta retrieved.fasta
~~~

### F Supplementary Experiment Analysis

#### F.1 Comparison of the Running time between RSA vs MSA

A severe speed bottleneck limits the use of previous MSA-based methods. In this part, we add analysis on database construction time as well as give details for inference time calculation. We calculate the total time used in each retrieval inference by summing: *alignment time* and *retrieval time*, as shown in Figure 10. Alignment time is the time used when finding MSA sequences through alignment and aligning found sequences with HHblits. Retrieval time is the time used during dense retrieval, including calculating the embedding of the query sequence with GPU. It is notable from the figure that alignment itself is a computationally costly procedure.

**Figure 10:**
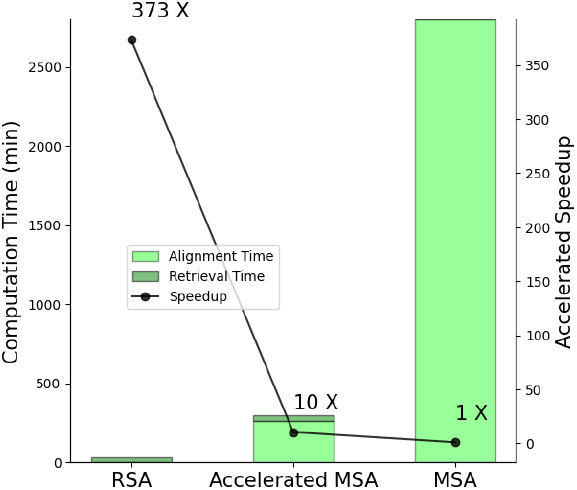
Illustration of speed up by RSA retrieval compared to MSA on secondary structure prediction dataset with 8678 sequences. Accelerated MSA refers to the MSA Transformer with MSA sequences retrieved by our RSA retriever.

Also, MSA is limited by its cubersome construction of retrieval HHM profile to perform HHM-HHM search. We follow the MSA custom database construction process in HHblits and compare with the construction time for RSA on a single V100 GPU (batch size=1) on a database of 10000 protein sequences. As shown in Figure 11, our method use only 10 minutes to finish the construction, though building a profile requires more than 3200 minutes.

**Figure 11:**
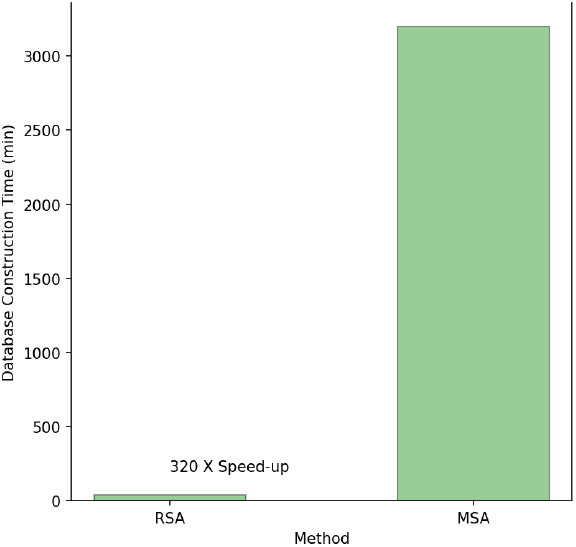
Illustration of speed up by RSA retrieval compared to MSA on database construction of 10000 protein sequences.

#### F.2 Case Study

We cherry-picked one example of ProtBERT and ProtBERT-RSA on homology prediction (1195 class classification task) to showcase the interpretability as well as give intuition on our method. As shown in Figure 12, our method takes the original sequence as well as 16 retrieved sequences for prediction. After weighted summing of all predicted results, the prediction of probability on ground truth label increase and gives the correct prediction. We checked the most highly weighted (top 5) retrieved sequences, all five proteins are Colicins, which is a family under Toxins’ membrane translocation domains. We can see from the case that weighting by distance helps the model focus on more similar retrieved instances.

**Figure 12:**
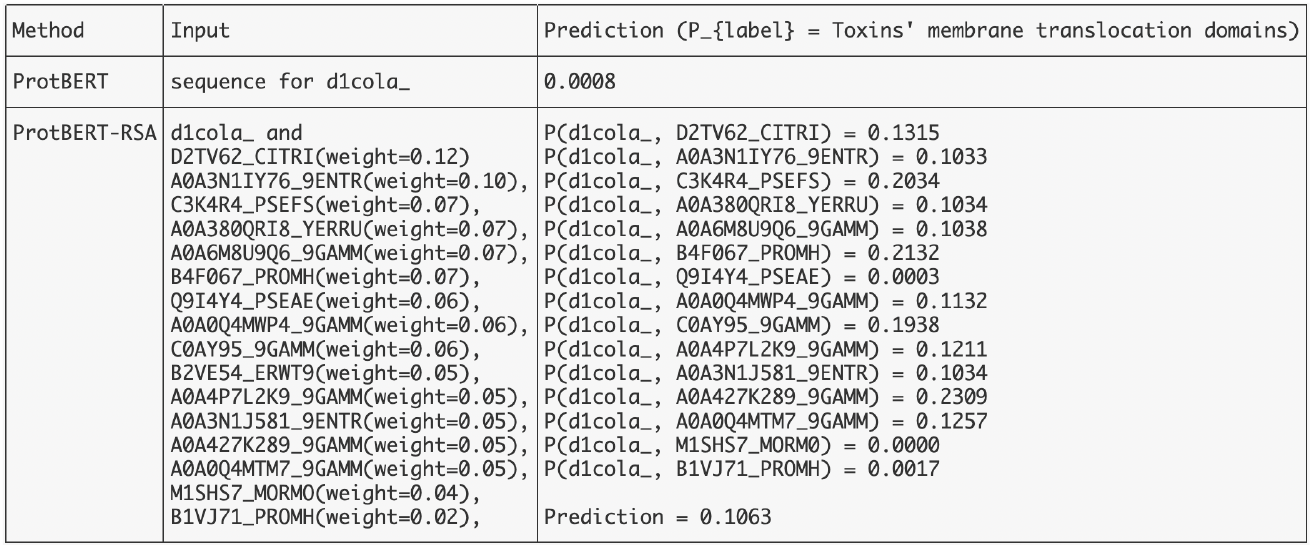
Case study on homology prediction.

**Figure 13:**
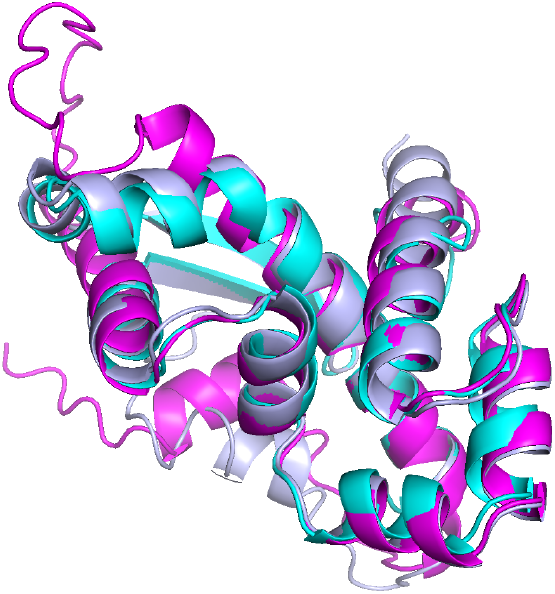
Structure Prediction for T1055, Cyan is the color for Ground truth. Pink is the color for ESMFold. Pink is the color for ESM-Fold. Light purple is the color for ESMFold-RSA. The TM-score for ESMFold is 0.70, and the TM-score for ESMFold-RSA is 0.91.

**Figure 14:**
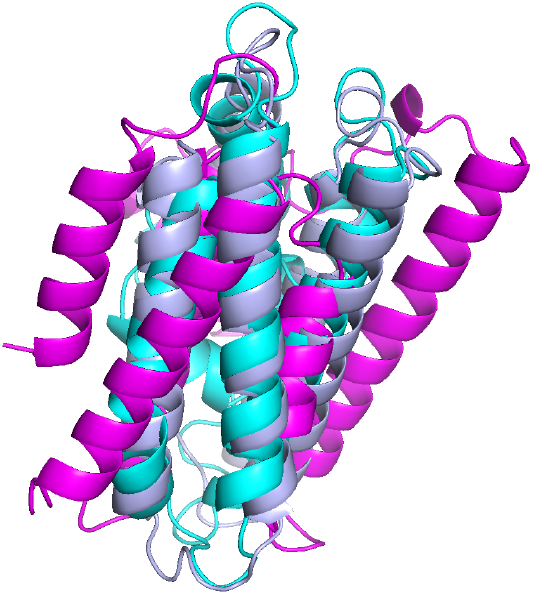
Structure Prediction for T1039, The TM-score for ESMFold is 0.61, and the TM-score for ESMFold-RSA is 0.29

We also provide two case studies on how RSA improves ESMFold. For target T1055, a DNA polymerase processivity factor, RSA retrieves *A0A1A8WBQ9_9APIC, A0A1Y4NGW6_9FIRM, A0A4V4NFM9_9ASCO, A0A1D3TXL7_9FIRM, A0A0V0QX86_PSEPJ, A9KN76_LACP7, A0A162CB07_9CRUS, A0A369KX60_9PROT,SKI2_SCHPO*, and the highest ranking augmentation prediction is from **(T1055, A0A1A8WBQ9_9APIC)**. A0A1A8WBQ9_9APIC is a Merozoite surface protein. Merozoite surface protein 7 (MSP7) is a protein of the malaria parasite that has been found to be associated with processed fragments from the MSP1 protein in a complex involved in red blood cell invasion. A0A1A8WBQ9_9APIC is a Merozoite surface protein C-terminal domain-containing protein that is related to DNA polymerase processivity factor through its requirement of a host factor, E. coli thioredoxin, in order to carry out its function. They also show similar structures with a TM-score of 0.42.

For target T1039, a virion RNA polymerase of crAss-like phage, RSA retrieves *A0A078ATM6_STYLE, A0A1D8P931_9FLAO, A0A363CW97_9PROT, D7JGI7_9BACT, A0A0B3VPN2_9FIRM, A0A1E4TQ27_PACTA, A0A1M6KY55_9FLAO, A0A1X7R9D3_9SACH, A0A0R1SCS6_9LACO, A0A367GMI1_9SPHI, A0A2N1F639_9FLAO, A0A0D6TLE8_9FLAO, A0A3N4NFZ1_9FLAO*,

*A0A1D2VEI9_9ASCO, A0A1L7I7H7_9FLAO, A0A1R0FA92_9RHIZ*. The highest ranking augmentation prediction is from **(T1039, A0A078ATM6_STYLE)**. A0A078ATM6_STYLE is a COMM domain-containing protein 1. It has no distinct functional relationship with T1039, though the second chain of this protein has a similar structure to T1039, with a TM-score of 0.34.

#### F.3 Domain Adaptation Analysis

In this section, we perform additional analysis on the domain adaptation ability on secondary structure prediction tasks. We perform training on NetSurfP-2.0[33] training set and test on two datasets with domain gaps. On CASP12, RSA marginally outperforms other baselines, as shown in Table 8. We also test on 10 de novo proteins (6YWC, 2LUF, 7BPM, 7BPL, 7CBC, 1FSD, 1IC9, 5JI4, 5KWO, 6W6X). Since we didn’t find secondary structure labels for these proteins, we provide visualization in Figure 15, which shows that our model has an obvious overhead over MSA Transformer on predicting geometric components.

**Table 8:** The performance of retrieval augmentation models w.r.t. the number of retrieved sequences on contact prediction.

| Methods | N=1 | N=4 | N=8 | N=16 | N=32 | N= full |
|---|---|---|---|---|---|---|
| Potts Model | — | 0.412 | 0.471 | 0.479 | 0.480 | 0.507 |
| MSA Transformer | 0.397 | 0.579 | 0.560 | 0.618 | 0.669 | — |
| Accelerated MSA Transformer | 0.397 | 0.524 | 0.538 | 0.608 | 0.654 | — |
| RSA | **0.556** | **0.595** | **0.615** | **0.717** | **0.719** | — |

**Table 9:** Overview for datasets in downstream tasks

| Task Name | Dataset source | #train sequences | #test sequences |
|---|---|---|---|
| Secondary Structure Prediction | NetSurfP-2.0 [33] | 8,678 | 513 |
| Contact Prediction | ProteinNet [3] | 25,299 | 40 |
| Remote Homology Prediction | Deepsf [25] | 12,312 | 718 |
| Stability Prediction | Rocklin's Dataset [46] | 53,571 | 12,851 |
| Subcellular Localization | DeepLoc [2] | 8,945 | 2,768 |
| Protein Protein Interaction | Pan's Dataset [39] | 6,844 | 227 |
| Protein Folding | CASP14 [34] | – | 65 |

**Table 10:** The domain adaptation performance of models on CASP12 secondary structure prediction.

| Method | CASP12 |
|---|---|
| ProtBERT | 0.628 |
| MSA Transformer | 0.621 |
| Accelerated MSA Transformer | 0.620 |
| RSA (ProtBERT backbone) | **0.631** |

**Figure 15:**
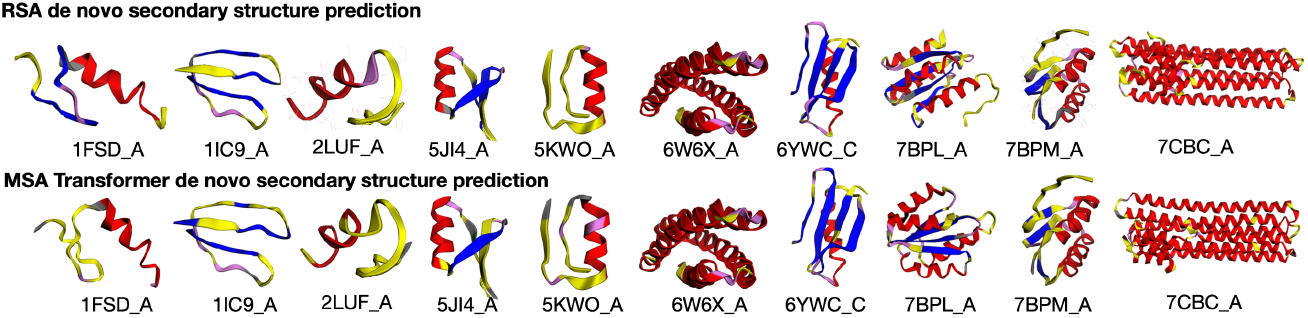
Prediction of Secondary Structure on De Novo Dataset. Each color corresponds to a different secondary structure.

#### F.4 Comparison of Accelerated MSA vs MSA quality

Accelerated MSA performs worse than original MSA when directly applied to MSA Transformer, as well as AlphaFold. In this section, we showcase successful and failed cases in AlphaFold and compare the coverage of two kinds of MSA.

As shown in Figure 18, AlphaFold prediction is closely correlated to the coverage of MSA sequence. On cases where dense retriever fails to find a wide coverage of homologous sequences, AlphaFold performances drop starkly. Note that the MSA is implemented as ColabFold [37], using Uniclust30 for MSA building, while our retriever database has a smaller coverage, using only Pfam database. Also we build accelerated MSA based on only top-500 sequences from retrieval.

#### F.5 Interpretability of RSA

In addition to analysis on interpretability in §5.7 in the main paper, we provide further analysis of the interpretability of RSA in terms of homology and structures.

##### Retrieval rank does not necessarily corresponds to the sequence closest to the query sequence token-wise

As shown in Figure 16, we calculate and rank the E-value of Top-32 retrieved protein sequences in CB513 dataset. We then calculate the average rank for the 1st, 2nd,… 32nd proteins in the dataset. It shows that the top-1 protein only has an average rank of 11, indicating that the retrieval rank does not necessarily corresponds to the sequence closest to the query sequence token-wise. Therefore, using dense retrieval, our retrieved results are diverse in sequences, though close to the query sequences in properties.

**Figure 16:**
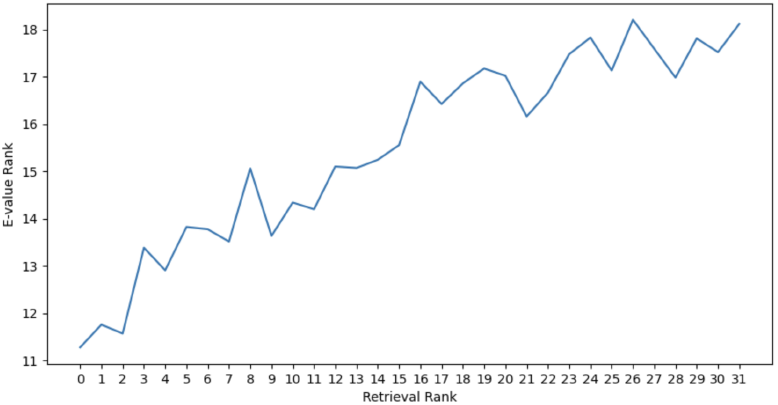
E-value rank against dense retrieval rank on in CB513 dataset.

##### Visualization on Retrieval Structural Similarity

As shown in Figure 17, we random picked a few more examples to illustrate the structural similarity between query protein and retrieval proteins. From the visualization, we can see that retrieved proteins exhibit similar structure or structure components, which could be used to boost structural knowledge.

**Figure 17:**
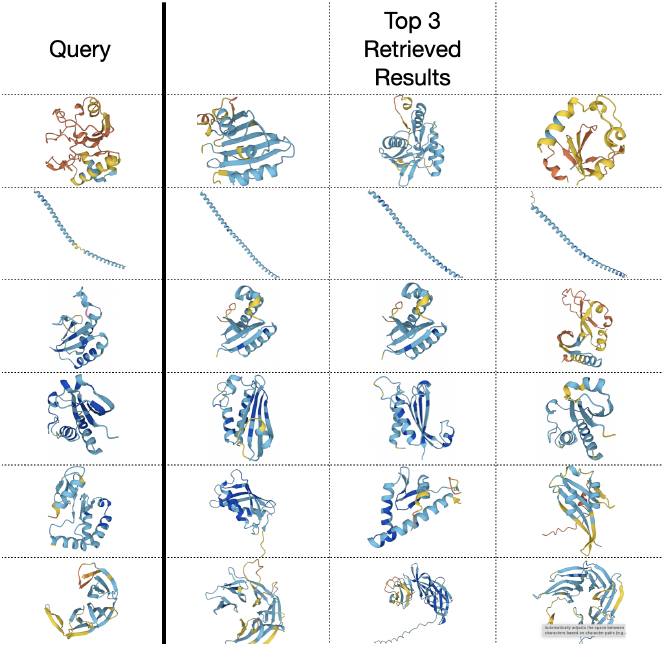
Query and Retrieved Sequence Structures

**Figure 18:**
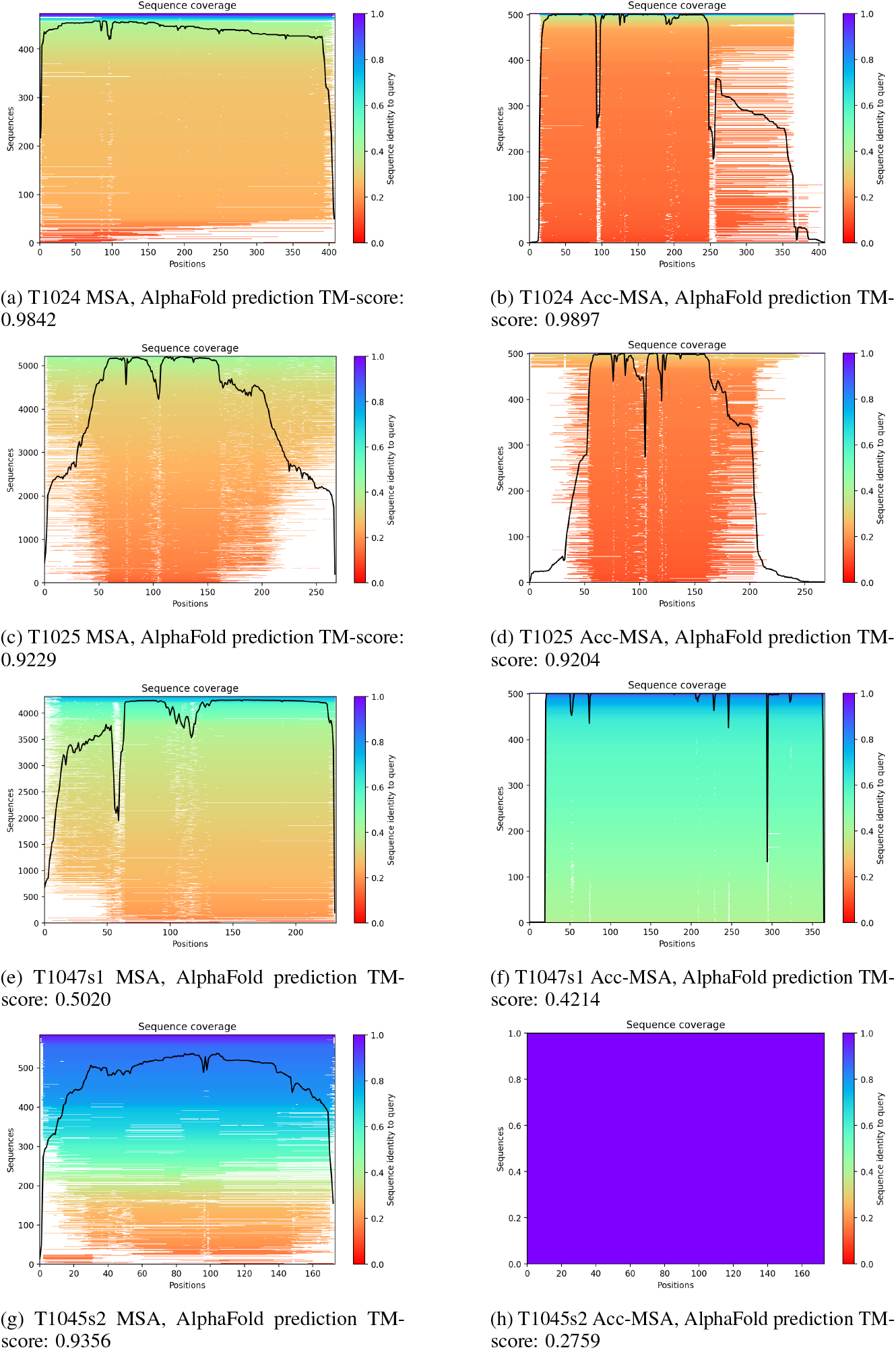
Visualization of the coverage rate of Accelerated MSA VS MSA.

#### F.6 ProteinChat: RSA Empowers ChatGPT on Protein Understanding

In the section, we point out a direction for our future work. RSA can not only be used on small-scale representation learning model, it can also augment large language models, e.g. ChatGPT. Currently, even GPT4 model shows limited understanding of biological sequences, especially long sequences like proteins. Galactica [49] pretrains LLM on scientific text and biological sequences, however, calibration on the nature of these sequences are ubiquitous. Here, we propose a new method to enable ChatGPT to understand protein sequences, **ProteinChat**, by directly applying the protein dense retriever to ChatGPT. We implement ProteinChat by binding ChatGPT with three tools,

- RSA retriever, that finds homologous sequences to the query protein.
- Annotator, that annotates the homologous sequences in a protein online database, i.e. InterPro^5^.
- Wikipedia Search, that facilitates more elaborated enquiry into the features of these proteins.

ProteinChat is implemented similar to ReAct agent [57], which views ChatGPT as an agent that uses chain-of-thoughts to break down a difficult task into parts that could be accomplished by available tools. Here we show several demos of ProteinChat.

Demo 1:

Question: Which family does this protein MSLNAEQKAKVVLEHGSSAHDTGSTEVQVAL LTLRINDLQKHFLEHKKDHHSRRGLLRMVSQRRKLLDYLKKRNISKYTDLIQSLGLRK

belongs to? (Ground truth: Belongs to the universal ribosomal protein uS15 family.) Thought: I need to find out the functional and evolutionary annotations of this protein. Action: Protein Retrieval

Action Input: MSLNAEQKAKVVLEHGSSAHDTGSTEVQVALLTLRINDLQKHFLEHKKD-HHSRRGLLRMVSQRRKLLDYLKKRNISKYTDLIQSLGLRK

Observation: Here are 4 homologs for this protein separated by “,”:B4D3S0_9BACT, RS15_HAMD5, K6ZQA3_9ALTE, A0A1H6FC85_9GAMM

Thought: I need to use the entry labels to search the InterPro database. Action: InterPro Annotation, Action Input: B4D3S0_9BACT

Observation: This is a 30S ribosomal protein S15. It is found in organism Chthoniobacter flavus Ellin428.It has molecular function that structural constituent of ribosome. It has a biological process translation. It has cellular component ribosome.

Thought: I need to search Wikipedia to find out which family this protein belongs to. Action: Wikipedia Search, Action Input: 30S ribosomal protein S15

Observation: Page: **Ribosomal protein** Summary: A ribosomal protein (r-protein or rProtein) is any of the proteins that, in conjunction with rRNA, make up the ribosomal subunits involved in the cellular process of translation. E. coli, other bacteria and Archaea have a 30S small subunit and a 50S large subunit, whereas humans and yeasts have a 40S small subunit and a 60S large subunit. **Page: 40S ribosomal protein S30** Summary: 40S ribosomal protein S30 is a protein that in humans is encoded by the FAU gene.

Thought: I now know the answer.

Final Answer: The protein belongs to the 40S ribosomal protein S30 family.

Demo 2:

Question: Can this protein MPGVTVKDVNQQEFVRALAAFLKKSGKLKVPEWVDTVK-LAKHKELAPYDENWFYTRAASTARHLYLRGGAGVGSMTKIYGGRQRNGVMPSHFSRGSKSVARRVLQALEGLKMVEKDQDGGRKLTPQGQRDLDRIAGQVAAANKK binds with

RNA? (Ground truth: Yes)

Thought:I need to find out what this protein does Action: Protein Retrieval

Action Input: MPGVTVKDVNQQEFVRALAAFLKKSGKLKVPEWVDTVKLAKHKE-LAPYDENWFYTRAASTARHLYLRGGAGVGSMTKIYGGRQRNGVMPSHFSRGSKSVARRVLQALEGLKMVEKDQDGGRKLTPQGQRDLDRIAGQVAAANKK

Observation: Here are 4 homologs for this protein separated by “,”:A0A315V8B8_GAMAF

,A0A673GQT0_9TELE, A0A2K6SCP9_SAIBB, A0A674GDP2_TAEGU

Thought: I need to find out what this protein does

Action: InterPro Search, Action Input: A0A315V8B8_GAMAF

Observation: This is a 40S ribosomal protein S19. It is found in organism Gambusia affinis (Western mosquitofish).

Thought: I need to find out if this protein binds with RNA

Action: Wikipedia Search, Action Input: 40S ribosomal protein S19 binds RNA

**Page: 40S ribosomal protein S19** Summary: 40S ribosomal protein S19 is a protein that in humans is encoded by the RPS19 gene. **Page: 5S ribosomal RNA** Summary: The 5S ribosomal RNA (5S rRNA) is an approximately 120 nucleotide-long ribosomal RNA molecule with a mass of 40 kDa. It is a structural and functional component of the large subunit of the ribosome in all domains of life (bacteria, archaea, and eukaryotes), with the exception of mitochondrial ribosomes of fungi and animals. The designation 5S refers to the molecule’s sedimentation velocity in an ultracentrifuge, which is measured in Svedberg units (S). **Page: Ribosomal protein** Summary: A ribosomal protein (r-protein or rProtein) is any of the proteins that, in conjunction with rRNA, make up the ribosomal subunits involved in the cellular process of translation. E. coli, other bacteria and Archaea have a 30S small subunit and a 50S large subunit, whereas humans and yeasts have a 40S small subunit and a 60S large subunit.)

Thought: I now know the final answer.

Final Answer: Yes, 40S ribosomal protein S19 binds with 5S ribosomal RNA as part of the ribosomal subunits involved in the cellular process of translation

## Footnotes

5 https://www.ebi.ac.uk/interpro/

## Notes

https://github.com/HKUNLP/RSA

### Competing Interest Statement

The authors have declared no competing interest.

### Summary of Updates

- Revised the main paper - Add protein folding experiments. - Provide a more detailed appendix.

